# Mitochondrial genes affect the immune response against bacterial infection (Duck Pasteurellosis) through a cascade of mechanisms mediated by nuclear immune genes-a cross talk with nuclear and mitochondrial gene

**DOI:** 10.1101/2023.08.30.555483

**Authors:** Argha Chakraborty, Aruna Pal, Manti Debnath, Jyoti Sahu, Paresh Nath Chatterjee, Abantika Pal, Joydeep Mukherjee, Rajarshi Samanta

## Abstract

Mitochondria possess 37 genes of its own, helpful for carrying out various functions including oxidative phosphorylation. But the mitochondria is neither independent in terms of its structure, nor its function, it depend upon nuclear gene for its functionality and replication. Our lab has earlier reported the role of mitochondrial genome in controlling health and reproduction in animal model. Reports also indicate that mitochondria have role in immune response against bacterial infection. In our current study, we depict the role of two polypeptide coding mitochondrial gene as cytochrome B and cytochrome C in providing host immunity against bacterial disease (Duck Pasteurellosis). We further observe that the mechanism was governed by cascade of mechanisms mediated through nuclear genes NRLP3, IL18 and Sting. In the first phase of the study, we have characterized Cytochrome B and Cytochrome C genes from Bengal duck and quantified through the assesment of the expression profiling with respect to healthy and Pasteurella infected ducks following a natural challenge with *Pasteurella multocida*. In the next phase of our study, we have characterized the above nuclear genes in Bengal duck and attempted to correlate the expression with mitochondrial genes. We attempted to explore the mechanism of pathway how mitochondrial DNA triggers the nuclear immune response leading to destruction of bacterial pathogen. Molecular docking revealed how IL18 directly binds with the pathogen. This is the first report in animal model.

## Introduction

Host immunity is important to understand Host-pathogen interaction, which is a dynamic process between diverse pathogens and host in all stages of pathogenic infection. Innate immune response is mediated through a cascade of mediators or proteins, each controlled by respective genes. We refer them as immune response genes, generally considered as nuclear genes regulating PRR, proinflammatory and anti inflammatory cytokines. In our lab,we have reported the role of certain set of nuclear genes in controlling disease resistance or animal health in sheep^1,2^, cattle^3^, buffalo^4^, goat^5,6^, chicken^7^ and duck^8,9^ model^10^. Simultaneously, we have also reported the role of mitochondrial genes in controlling health and disease resistance^11^, and even reproduction^12^. Mitochondrial genes not only help through oxidative phosphorylation, but it also help in other series of metabolic processes, including calcium^13^ and iron^14^ metabolic pathways. We have already reported certain preliminary works^15^ involving all the 37 mitochondrial genes in duck through next generation sequencing approach of Whole mitochondrial genome sequencing^16,17^. Apart from it, we had also studied duck genomics^18–20^. Now recent evidences indicate the role mitochondrial DNA in regulation of nuclear immune response genes and thereafter the pathogen is destroyed^21^.

Duck cholera, which is known to be caused by Pasteurella multocida, is known to result in high morbidities and fatalities as well as significant economic losses. Chronic duck cholera results in the localisation of the organism in specific organs and is distinguished from acute duck cholera by severe watery diarrhoea, anorexia, respiratory symptoms, and septicemia. An outbreak flock of 50,000 birds had an average cost of $0.40 (€0.33) per bird, while nonoutbreak flocks that had received the fowl cholera vaccine had an average cost of $0.12 (€0.10) per bird. Furthermore, the cost of treating an outbreak of fowl cholera in 1985–86 was estimated to be $0.51 (€0.452) per bird^22^. This cost included productivity losses as well. About half as much was spent on the immunisation against poultry cholera overall as there was on the outbreak. In all, California lost 0.5% of its whole 1985–1986 turkey meat supply due to fowl cholera.

Ducks have been considered to be the second most important poultry species after chicken. Indigenous ducks have been observed to have certain unique characteristics in terms of its disease resistance ability against common avian diseases^8^. Ducks can be easily reared under organized farming system with small water container that enable them to dip their head for washing of their eyes without any natural water bodies. It is one of the least explored species with certain unique characteristics in response to innate immune response.

Hence in this current study, we attempt to characterize certain nuclear innate immune response genes and polypeptide coding mitochondrial genes and to explore their role and certain interaction among them in providing immune response against duck Pasteurellosis (bacterial disease) in duck model.

## Materials and Methods

### I. Characterization of Mitochondrial genes (Cytochrome B & Cytochrome C) and nuclear gene (NRLP3, IL18 and Sting) in liver tissue of Bengal Duck

#### Birds and sample collection

The samples were collected from Bengal ducks maintained at the West Bengal University of Animal and Fishery Sciences.

#### Sample Collection and RNA Isolation

Duck liver tissue (1g) was collected from slaughtered birds which were clinically healthy and maintained in the duck house of the LFC Dept, WBUAFS. Adult males (n=6) were selected for the collection of samples. The tissue was immersed in Trizol in the vial and transported in ice to the laboratory for RNA isolation. Total RNA was isolated using the TRIzol extraction method (Life Technologies, USA), as per the standard procedure and was further utilized for cDNA synthesis^23–26^. cDNA concentration was estimated and samples above 1200 micrograms per ml were considered for further study. Tissues from gut-associated lymphoid tissue (GALT) as ileocaecal junction, liver, gizzard, caecum, jejunum, spleen, and pancreas were collected from both healthy and duck infected with duck plague virus for studying the expression profiling with quantitative PCR.

#### cDNA synthesis and PCR Amplification of the Mitochondrial genes (Cytochrome B & Cytochrome C) and nuclear gene (NRLP3, IL18 and Sting) gene of Bengal Duck

20μL of the reaction mixture was composed of 5μg of total RNA, 0.5μg of oligo dT primer (16–18mer), 40U of Ribonuclease inhibitor, 1000M of dNTP mix, 10mM of DTT, and 5U of MuMLV reverse transcriptase in an appropriate buffer. The reaction mixture was mixed thoroughly followed by incubation at 37°C for 1 hour. The reaction was allowed up to 10 minutes by heating the mixture at 70°C unliganded and then chilled on ice. Afterwards, the integrity of the cDNA was checked by performing PCR^23–26^. The concentration of cDNA was estimated through Nanodrop. **NRLP3, IL18, Sting, Cytochrome B and Cytochrome C** primer pairs were designed based on the published sequences of chicken and duck origin using DNASTAR software (Hitachi Miraibio Inc., USA) to amplify full-length open reading frame (ORF) of these genes (Table 1).

**Table 1a.**
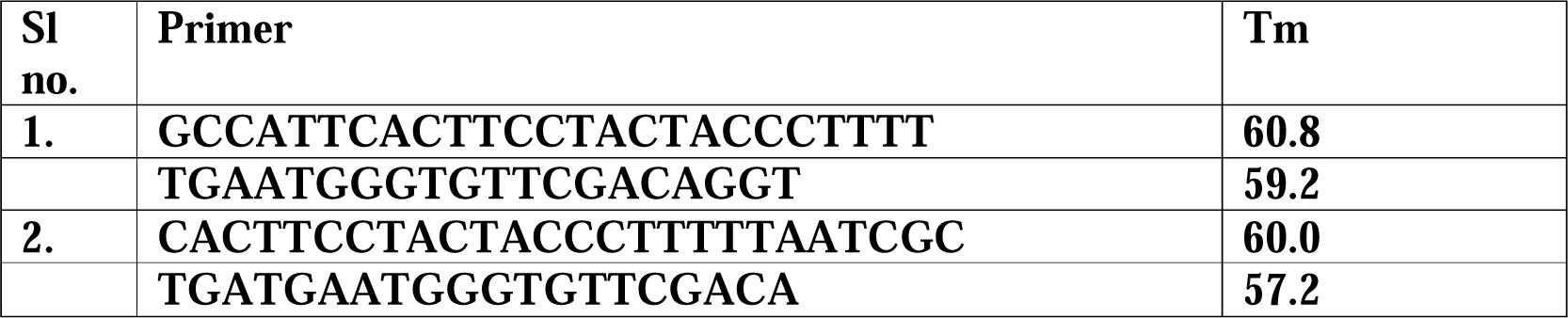
Primer for Mitochondrial genes (Cytochrome B) of Bengal Duck.

**Table 1b:**
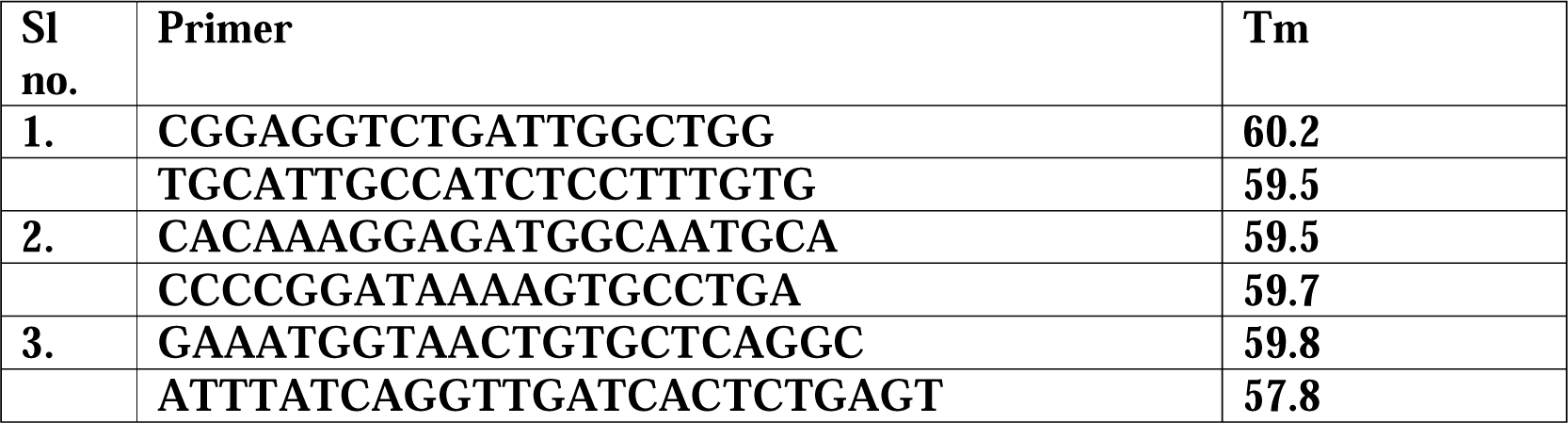
Primers for Mitochondrial genes (Cytochrome C) in liver tissue of Bengal Duck.

**Table 1c.**
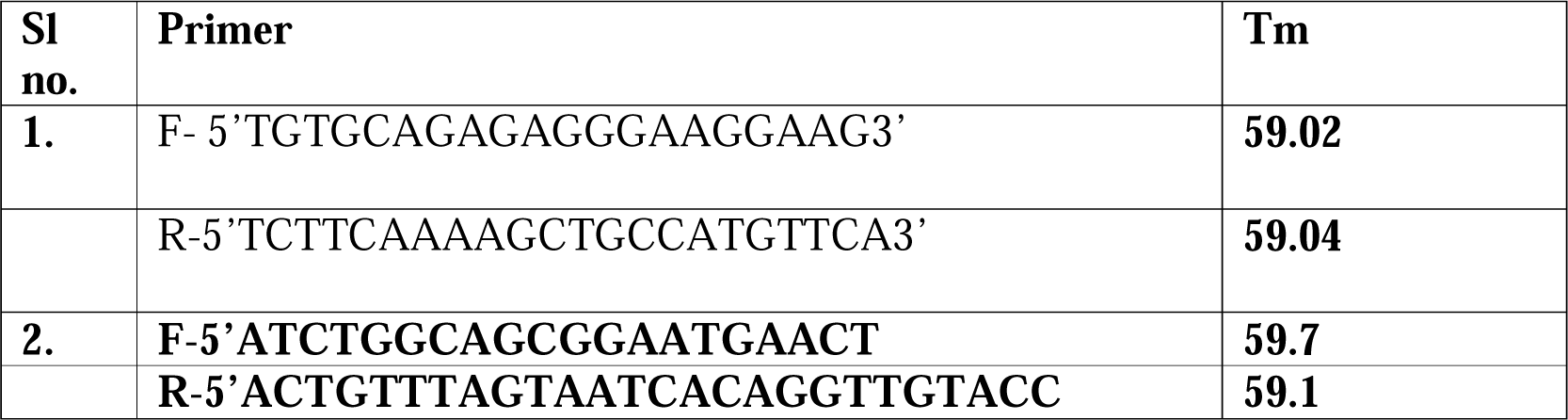
Primer for amplification of IL18 gene in Duck.

**Table 1d.**
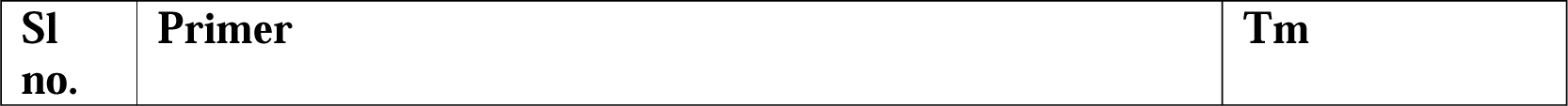
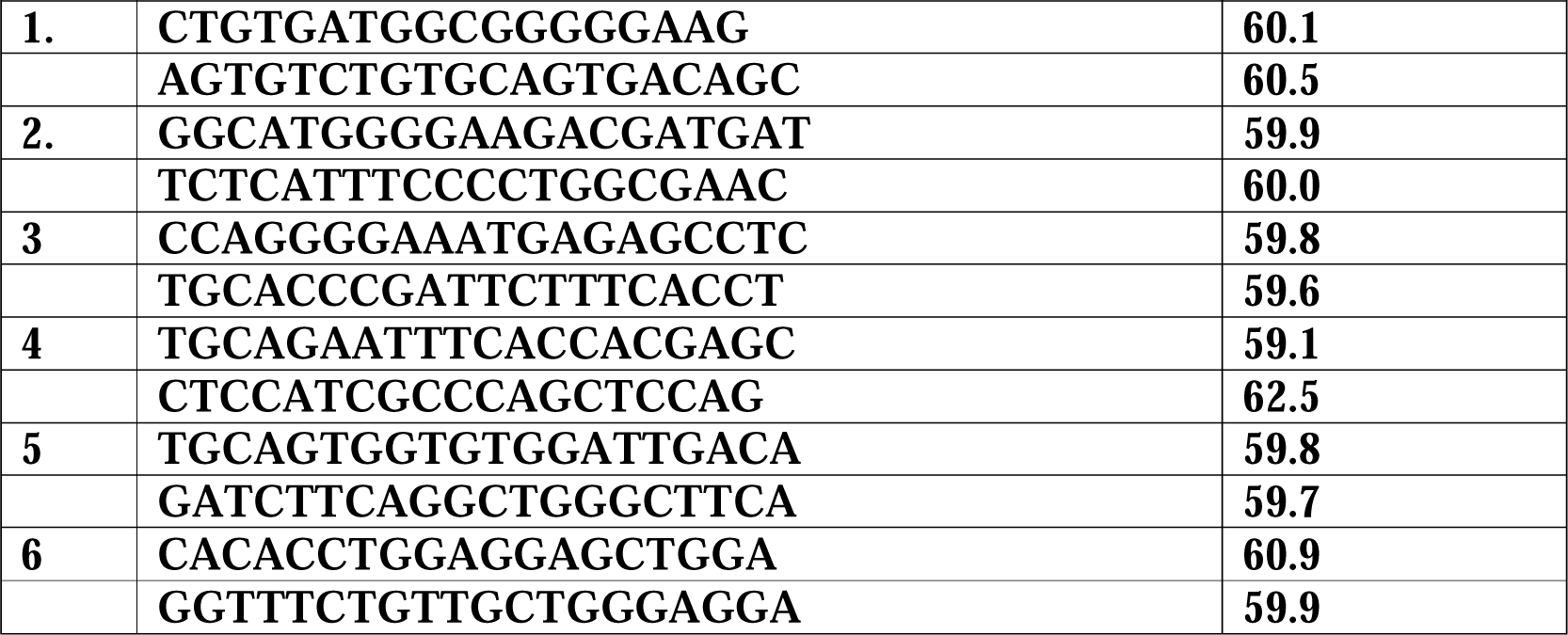
Primer for amplification of NRLP3 gene in duck.

**Table 1e.**
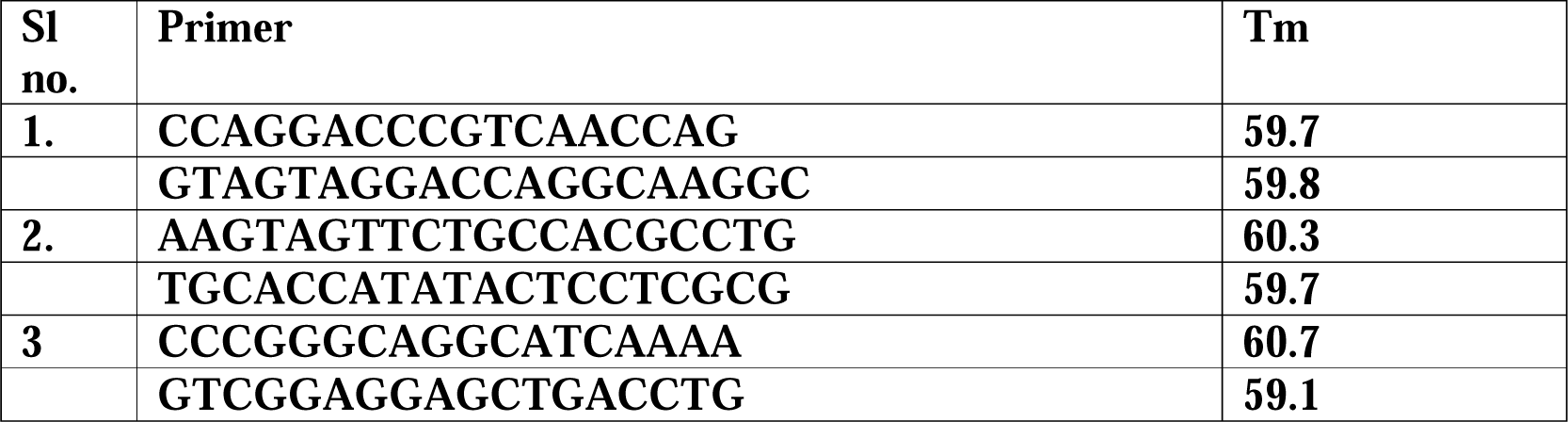
Primer for amplification of Sting gene in duck.

**Table 2:**
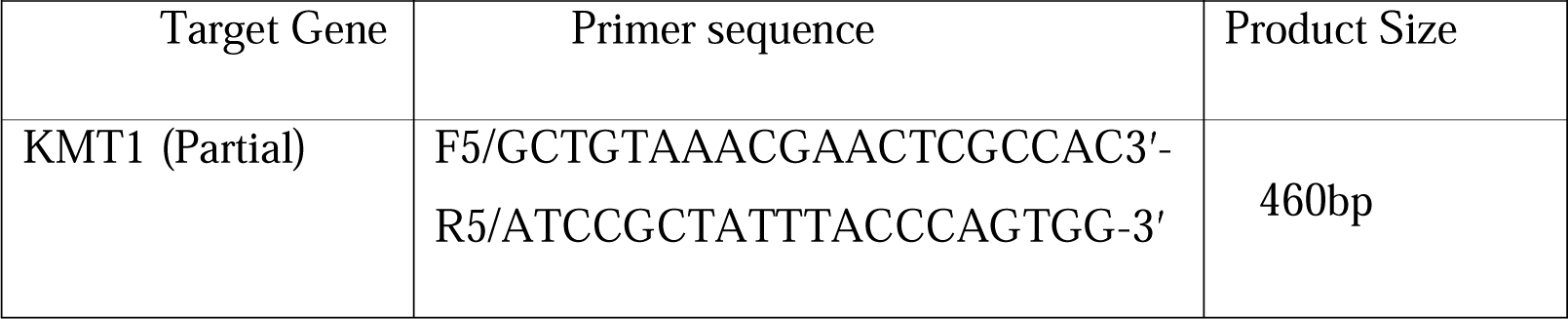
Primers used for the molecular detection of Pasteurella multocida.

**Table 3:**
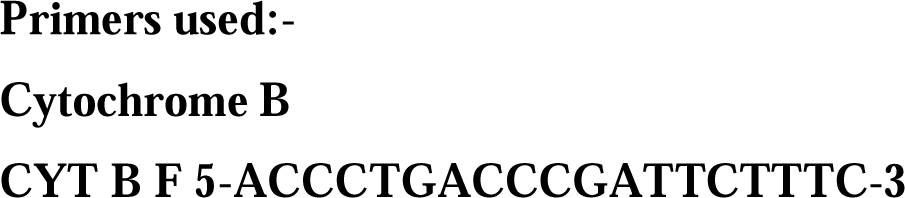
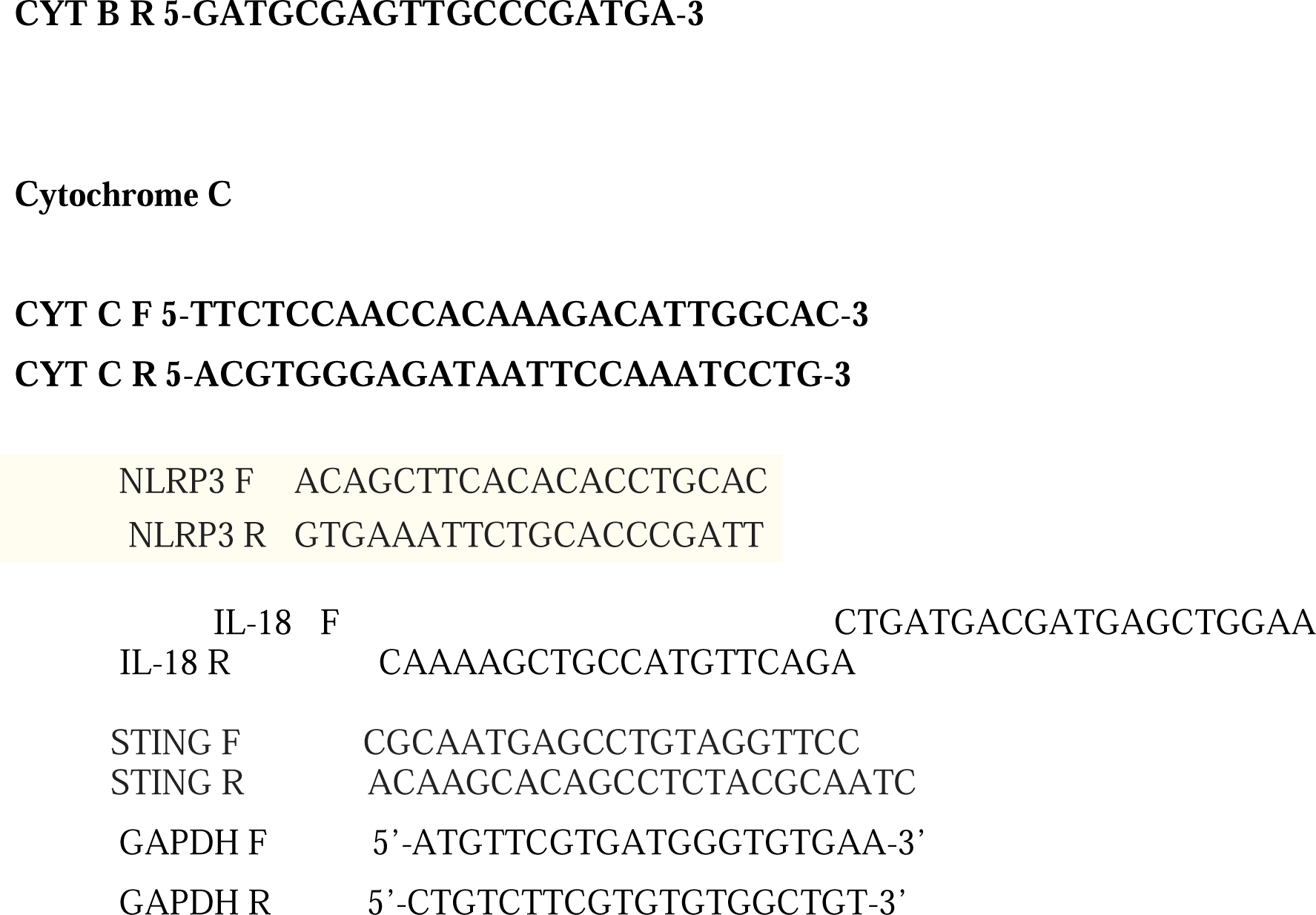
Primers used for the differential mRNA expression profiling of immune response genes of duck.

#### Mitochondrial genes (Cytochrome B & Cytochrome C) and nuclear gene (NRLP3, IL18 and Sting)

25μL of reaction mixture was comprised of 80–100ng cDNA, 3.0μL 10X PCR assay buffer, 0.5μL of 10mM dNTP, 1U Taq DNA polymerase, 60ng of each primer, and 2mM MgCl2. PCR reactions were performed in a thermocycler (PTC-200, MJ Research, USA) with cycling conditions as, initial denaturation at 94°C for 3min, further denaturation at 94°C for 30sec, annealing at 61°C for 35sec, and extension at 72°C for 3min were conducted for 35 cycles followed by a final extension at 72°C for 10 min. The list of primers pertaining to these genes are listed in Table 1 (a-e).

**Characterization of nuclear genes (NP, IL18 and String) in Bengal Duck**

### cDNA Cloning and Sequencing

Amplified products of RIGI, MDA5 and interferon-alpha were checked by 1% agarose gel electrophoresis. The products were purified from gel using a Gel extraction kit (Qiagen GmbH, Hilden, Germany) .pGEM-T easy cloning vector (Promega, Madison, WI, USA) was used for cloning. Then, 10μL of the ligated product was mixed thoroughly with 200μL competent cells, and heat shock was given at 42°C for 45sec in a water bath. Subsequently, the cells were immediately transferred on chilled ice for 5 min., and SOC medium was added to it. The bacterial culture was centrifuged to obtain the pellet and plated on an LB agar plate containing Ampicillin (100mg/mL) added to the agar plate @1:1000, IPTG (200mg/mL) and X-Gal (20mg/mL) for blue-white screening. Plasmid isolation from overnight-grown culture was carried out by a small-scale alkaline lysis method as described^21^. Recombinant plasmids were characterized by PCR using CD14 primers as reported earlier and restriction enzyme digestion. CD14 gene fragments released by enzyme EcoRI (MBI Fermentas, USA) were inserted in a recombinant plasmid which was sequenced by the dideoxy chain termination method with T7 and SP6 primers in an automated sequencer (ABI prism, Chromous Biotech, Bangalore).

### Sequence Analysis

DNASTAR Version 4.0, Inc., USA software was employed for the nucleotide sequence analysis for protein translation, sequence alignments, and contigs comparisons. Novel sequences were submitted to the NCBI Genbank and accession numbers were obtained which are available in the public domain.

### Study of Predicted duck Mitochondrial (Cytochrome B & Cytochrome C) and nuclear (NRLP3, IL18 and Sting) Protein Using Bioinformatics Tools

Predicted peptide sequences of duck **Mitochondrial (Cytochrome B & Cytochrome C) and nuclear (NRLP3, IL18 and Sting)** were deduced. Protein were then aligned with that of other species using MAFFT^27^. The analysis was conducted for a sequence-based comparative study. The signal peptide is essential to prompt a cell to translocate the protein, usually to the cellular membrane and ultimately signal peptide is cleaved to give a mature protein. Prediction of the presence and location of the signal peptide of these genes was conducted using the software (SignalP 3.0 Sewer-prediction results, Technical University of Denmark). The leucine percentage was calculated manually from the predicted peptide sequence. Di-sulphide bonds are essential for protein folding and stability, ultimately. It is the 3D structure of the protein which is biologically active.

Protein sequence level analysis was employed (http://www.expasy.org./tools/blast/) for the assessment of leucine-rich repeats (LRR), leucine zipper, detection of Leucine-rich nuclear export signals (NES), and detection of the position of GPI anchor, N-linked glycosylation sites. Since these are receptors, they are rich in leucine-rich repeats, which are essential for pathogen recognition and binding. A leucine zipper is essential to assess the dimerization of IR molecules. N-linked glycosylation is important for the molecule to determine its membranous or soluble form. The leucine-rich nuclear export signal is essential for the export of this protein from the nucleus to the cytoplasm, whereas GPI anchor is responsible for anchoring in the case of membranous protein. Prosite was used for LRR site detection.

Leucine-rich nuclear export signals (NES) were analyzed with NetNES 1.1 Server, Technical University of Denmark. O-linked glycosylation sites were detected using NetOGlyc 3.1 server (http://www.expassy.org/), whereas N-linked glycosylation sites were assessed through NetNGlyc 1.0 software (http://www.expassy.org/). Sites for leucine-zipper were detected through Expassy software, Technical University of Denmark^28^. Sites for alpha helix and beta sheet were detected using NetSurfP-Protein Surface Accessibility and Secondary Structure Predictions, Technical University of Denmark^29^. Domain linker sites were predicted^30^. LPS-binding^31^ and LPS-signalling sites^32^ were predicted based on homology studies with other species of respective polypeptide. These sites are important for pathogen recognition and binding.

### Three-dimensional structure prediction and Model quality assessment

Three-dimensional models of **Mitochondrial protein (Cytochrome B & Cytochrome C) and nuclear protein (NRLP3, IL18 and Sting)** polypeptide were predicted through the Swissmodel repository^33^. Templates possessing the greatest identity of sequences with our target template were identified with PSI-BLAST (http://blast.ncbi.nlm.nih.gov/Blast). PHYRE2 server based on the ‘Homology modelling approach was used to build three dimensional model of these proteins^34^. Molecular visualization tool as PyMOL (http://www.pymol.org/) was employed for model generation and visualization of the three-dimensional structure of these proteins understudies for duck origin. The structure of duck molecules was evaluated and assessed for its stereochemical quality (through SAVES, Structural Analysis and Verification Server, http://nihserver.mbi.ucla.edu/SAVES/); then refined and validated through ProSA (Protein Structure Analysis) web server (https://prosa.services.came.sbg.ac.at/prosa)^35^. NetSurfP server (http://www.cbs.dtu.dk/services/NetSurfP/ Peterson et al., 2009) was used for assessing the surface area of these proteins through relative surface accessibility, Z-fit score, and probability of alpha-Helix, beta-strand and coil score.

The alignment of 3-D structure of these proteins was analyzed with RMSD estimation to evaluate the structural differentiation by TM-align software^36^.

### Protein-protein interaction network depiction

To understand the protein interaction network of these proteins, we performed the search in STRING 9.1 database^37^. The functional interaction was assessed with a confidence score. Interactions with scores < 0.3, scores ranging from 0.3 to 0.7, and scores >0.7 are classified as low, medium and high confidence respectively. Also, we executed a KEGG analysis which depicted the functional association of these proteins with other related proteins.

### In silico study for the detection of the binding site of IL18 with Pasteurella multocida

#### Molecular Docking of host IL18 with the Pasteurella multocida surface proteins

Molecular docking is a bioinformatics tool used for in silico analysis for the prediction of the binding mode of a ligand with a protein 3D structure. Patch dock is an algorithm for molecular docking based on the shape complementarity principle^38^. Patch Dock algorithm was used to predict ligand-protein docking for surface antigen for *Pasteurella multocida* with the host immune response gene IL18. Firedock analysis was assessed for further confirmation.

### Sample Collection

For the present study, six tissue samples were collected from the infected indigenous duck samples followed by a challenge study and six tissue samples from six healthy duck following infection. We designate the ducks with mortality as infected or diseased (less immune) and the ducks that survived as healthy (better immune).

#### Natural Challenge of ducks with Pasteurella multocida, symptomatic diagnosis, PCR based detection

Bengal ducks in the experimental duck Farm were naturally infected with ***Pasteurella multocida*** following a natural disaster. We have grouped the ducks in two groups-Infected (with symptoms that ultimately die) and healthy (survivors without any symptom).

#### Challenge study with Pasteurella multocida in ovo, estimation of infective bacterial load at different stages of infection, PCR based detection

For the challenge study, initially, we incubated the fertile duck eggs (n=12) in the incubator at 37^0^C along with water in a Petri dish to maintain the proper humidity, which is essential for the incubation of duck eggs. After one week, we rechecked and screened the fertile embryo through candling. To harvest the **Pasteurella multocida bacterial culture**, at nine days of incubation of the fertilized egg, we puncture the air sac with the help of a sterilized needle through the chorioallantoic membrane and inoculated 200ul of the infective agent in a sterile environment. The needle puncture was quickly sealed with glue (feviquick). The eggs were again incubated in a sterile environment at 37^0^C and humid conditions as earlier. The entire procedure was undertaken at the BSL3 lab. The duck pasteurella bacterial strain employed was the available infection in the state from the clinical infected cases. After 3 days of inoculation, the bacteria were isolated from embryonic fibroblast cells through a viral DNA isolation kit. Quantification of bacterial load was estimated through quantitative PCR. To challenge, 0.3ml of bacterial stock at EID50 (Egg infective dose 50) was employed for each embryonated egg. Infective amnio-allantoic fluid was used to determine 50 per cent EID. Initial verification was conducted to assure that the ducks were free from Duck Pasteurella infection.

### Sample Collection

For the present study, six tissue samples were collected from the infected indigenous duck samples followed by a challenge study. We designate the ducks with mortality as infected or diseased (less immune) and the ducks that survived as healthy (better immune).

### Diagnosis and confirmation of Pasteurella multocida

The ducks were initially diagnosed with duck pasteurella clinically with the help of initial symptoms. Later on, the confirmatory diagnosis was conducted through molecular detection of the Duck pasteurella multocida specific primers in the infected samples.

### Gross Anatomical view of the liver in Pasteurella infected duck

We studied different body organs from infected ducks for clinical diagnosis of duck pasteurella.

### Confirmation through molecular detection

In the next step, we follow the confirmation of the samples with a molecular PCR-based technique using the following primers as recommended by OIE, Paris.

It includes bacterial DNA isolation through an isolation kit and is subjected to PCR.

### Assessment of dose of infection

We estimated the dose of bacterial infection standardization through Quantitative PCR. We take the standard vaccine (against duck cholera) as a control for its amplification and quantification.

### Differential mRNA expression profiling for Cytochrome B and Cytochrome C, NRLP3, IL18 and Sting in Bengal Duck

#### RT-PCR (Real Time-Polymerase Chain Reaction)

#### RNA Estimation

The tissue samples were cut into small pieces and submerged in liquid Trizol followed by thorough grinding with a mortar and pestle. 1 ml triturated tissue was taken into a 2 ml Eppendorf tube and 1 ml chloroform was added. Then it was centrifuged at the rate of 10,000 rpm for 10 minutes at 4°C in an automated temperature-controlled refrigerated centrifuge machine (BR Biochem, Life Sciences and REMI C24 plus). Three phases of differentiation were identified. The uppermost aqueous phase was collected for RNA isolation into a new Eppendorf tube and an equal volume of 100 percent isopropanol was added. It was left for a minute at 20-25°C temperature and centrifuged at the rate of 10,000 rpm for 10 minutes. After that supernatant was discarded and an equal volume of 70% ethanol was added to the pellet. Again the mixture was centrifuged at the rate of 10,000 rpm for 10 minutes and then the supernatant was discarded. Then the Eppendorf tube containing the pellet was kept at room temperature for air drying. After drying the remaining pellet dissolved with nuclease-free water (Applied Bio-systems, Cat. No. AM9930) and stored at - 20°C for future uses.

### Qualitative Analysis of Total RNA

The total RNA was quantified using NanoDrop-8000 (Thermo Scientific, Model No. 8000 Spectrophotometer), taking 1µl of each sample to determine the concentration and A260/280 ratio.

### First Strand cDNA Synthesis (Reverse Transcriptase-PCR)

5µl estimated RNA was taken into a PCR micro tube and added 1 µl OligoDT primer, 1µl 10mM dNTP and 13µl distilled water. After well mixing, it was heated at 65°C for 5 minutes. Then those tubes were quickly chilled on ice for 5-7 minutes and centrifuged. Then 4 µl 5x first strand buffer and 2 µl DTT (100mM) were added to the tube. It was well mixed and incubated at 37°C for 2 minutes. Then 1 µl M-MLVRT (200 u/µl) was added to the reaction mixture and treated at 37°C for 50 minutes and at 70°C for 15 minutes for inactivation. All the temperature treatments were maintained automatically in the thermal cycler machine (Applied Biosystems by Life Technologies) by setting the whole programme.

### Primer Designing and Synthesis

All the primers were designed using primer 3 software (v. 0. 4.0) as per the recommended criteria. MDA5, RIG1 and GAPDH used in the experiment, were designed (DNASTAR software).

### Real-Time PCR Experiment (SYBR Green based)

The primers were standardised with respective cDNA samples before being subjected to real-time PCR. The entire reactions were performed in triplicate (as per MIQE Guidelines) and the total volume of the reaction mixture was set up to 20µl. The reaction mixture is set up with MDA5, RIG1 and GAPDH (housekeeping gene) primers calculated as per concentration. 1µl of cDNA, 10µl Hi-SYBr Master Mix (HIMEDIA MBT074) and rest volume adjusted with nuclease-free water to achieve the total reaction mixture volume. Then the reaction plate placed into the ABI 7500 system and run the reaction program. The delta-delta-Ct (ΔΔCt) method 2^-ΔΔCt^ was used for the analysis of the result. The primers used for the reaction are as followed:

### Estimation of haematological and biochemical parameters

#### Haematological Profiles

The haematological parameters like haemoglobin, erythrocyte sedimentation rate (ESR) and packed cell volume (PCV) were estimated in whole blood soon after the collection of blood. Haemoglobin was estimated by acid haematin method^39^, E.S.R. and PCV by Wintrobe’s tube^40^. The total erythrocyte count (TEC), total leukocyte count (TLC) and Differential leukocyte count (DLC) were studied by standard methods described^41^.

### Biochemical Analysis

The serum biochemical parameters, estimated in the experiments were total protein, albumin, globulin, albumin: globulin, aspartate aminotransferase (AST), alanine aminotransferase (ALT), alkaline phosphatase (ALP), Total bilirubin, Indirect bilirubin, direct bilirubin, glucose, uric acid, urea and BUN by using a semi-auto biochemistry analyzer (Span diagnostic Ltd.) with standard kits (Trans Asia Bio-Medicals Ltd., Solan, HP, India). The methodology used for the estimation of total protein, albumin, total & direct bilirubin, ALT, ALP, glucose, creatinine urea and uric acid were the biuret method, bromocresol green (BCG) method, 2-4-DNPH method, modified kind and king’s method, GOD/POD method, modified Jaffe’s Kinetic method. GLDH-urease method and trinder peroxidise method respectively.

### Histological section

The liver samples were fixed in formalin (10%) and embedded in paraffin and processed for histological examination and stereology. The liver tissues were submerged in Lillie fixative for 1 week at room temperature and then were processed and embedded vertically in paraffin wax. Then, each liver sample was exhaustively sectioned into 4 μm-thick sections by a fully automated rotary microtome (Leica RM2255, Germany). Each of these sections was stained with hematoxylin and eosin and mounted. From each liver sample, 10-15 sections were chosen by the systematic uniform random sampling (SURS) method.

### Immunohistochemistry

Paraffin tissue blocks were prepared by standard manual alcohol-acetone protocol. The 5-6 µm thick paraffin sections obtained from the liver of both the infected and healthy control ducks, were taken on Millennia 2.0 adhesion slides (Cat. No. 71863-01, Abcam). The tissue sections were de-paraffinized and hydrated in distilled water. The tissue slides were covered with trypsin enzymatic antigen retrieval solution (Cat. No. ab970, Abcam) and kept in an incubator in a humid environment at 37°C for 5-10 minutes. The sections were then incubated for 60 minutes in peroxidase blocking solution (Lot. No. 00065614, Dako) at room temperature to block non-specific antibody binding activity. After subsequent washing with phosphate buffer saline (PBS), the sections were incubated at 37°C for 2 hours in a humid environment with mouse monoclonal anti-IL18 antibody in 1: 200 dilution. Immunoreactivity was detected after one-hour incubation at 37°C with a secondary antibody, Rabbit anti-mouse IgG H&L (HRP Conjugated, Cat. No. ab6728; Abcam) in dilution 1:200. Slides were then rinsed 3 times in PBS for 5 minutes each, followed by treatment with freshly prepared DAB solution for 3 minutes (DAB substrate, Cat. No. 34001, Thermo Fisher Scientific). The sections were counter-stained with Mayer’s haematoxylin, hydrated in ethanol, cleared in xylene and then mounted in DPX.

### Statistical Analysis

Microsoft Excel was used for the descriptive statistical analysis. SYSTAT 13.1 software (SYSTAT Software Inc.) was used for statistical analysis and analysis of variance (ANOVA) was used to test between parameters.

## Result

### Characterization of the Cytochrome B and Cytochrome C derived peptide from Duck liver

We have sequenced Cytochrome B gene for Bengal duck and predicted its amino acid sequence through in silico methods.In the next step, we have predicted the 3D model for Cytochrome B (Fig 1) and Cytochrome C (Fig 2) for Bengal duck.

**Fig 1:**
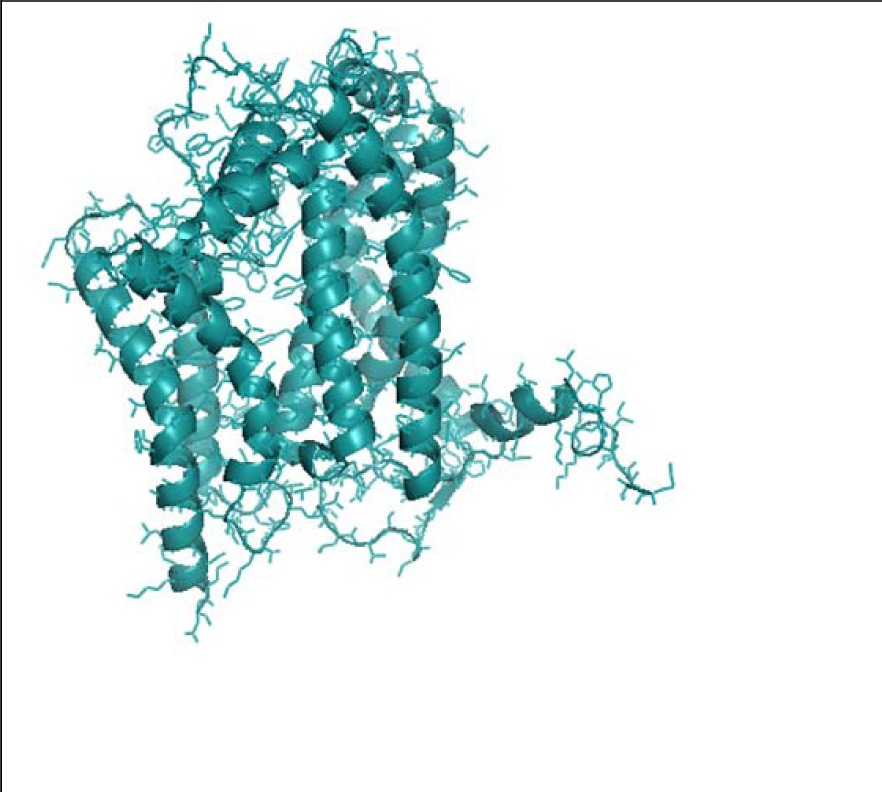
3D model for duck Cytochrome B.

**Fig 2:**
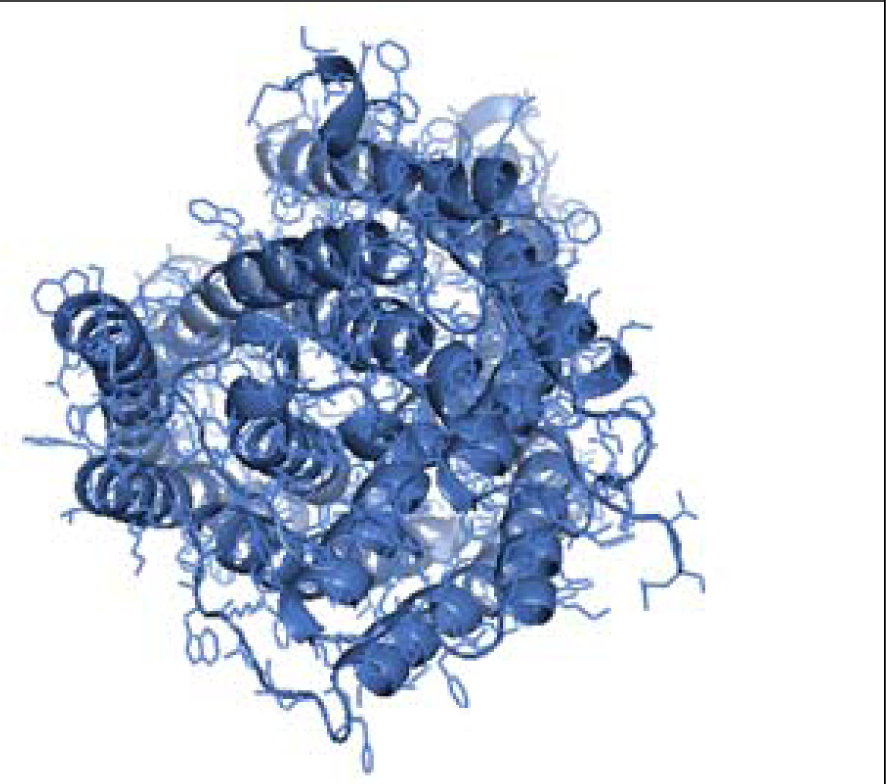
3D model for duck Cytochrome C.

We have obtained the 3D structure for NRLP3 (Fig 3), IL18 (Fig 4), and Sting (Fig 5) gene of Bengal Duck.

**Fig 3:**
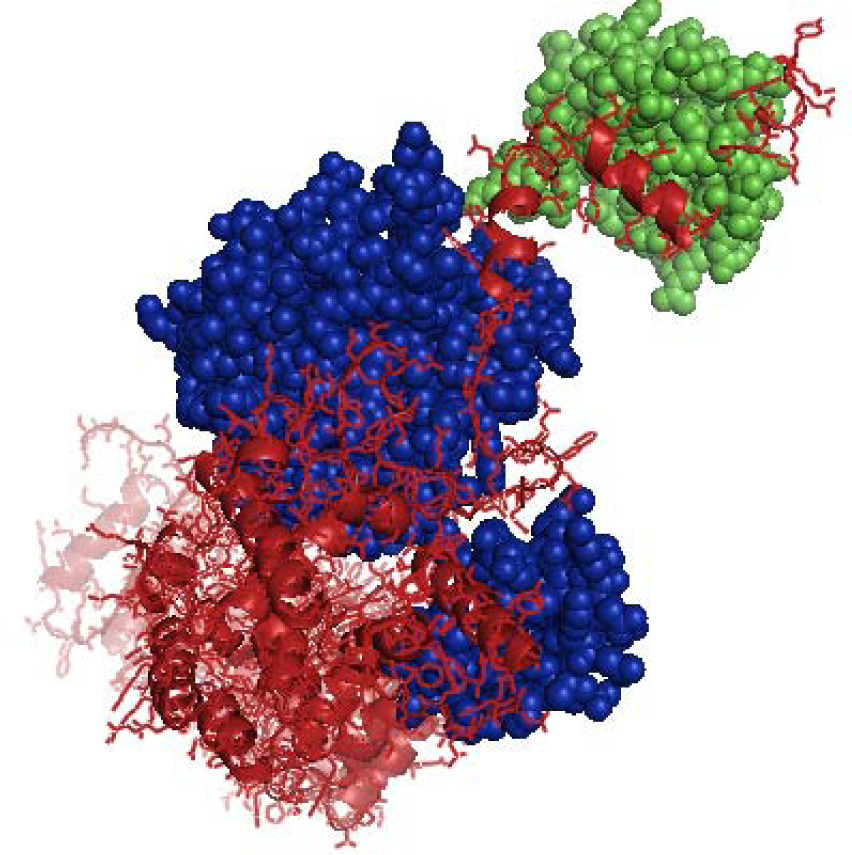
3D structure of NLRP3 green sphere: 1-94:DAPIN or Pyrin domain Blue sphere: 177-380: NACHT domain.

**Fig 4:**
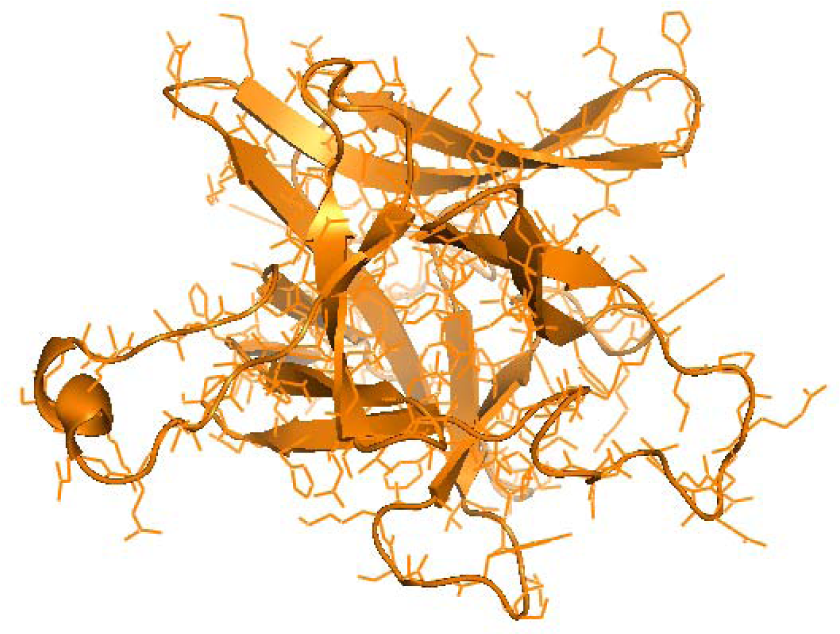
3D structure for IL18 for duck.

**Fig 5:**
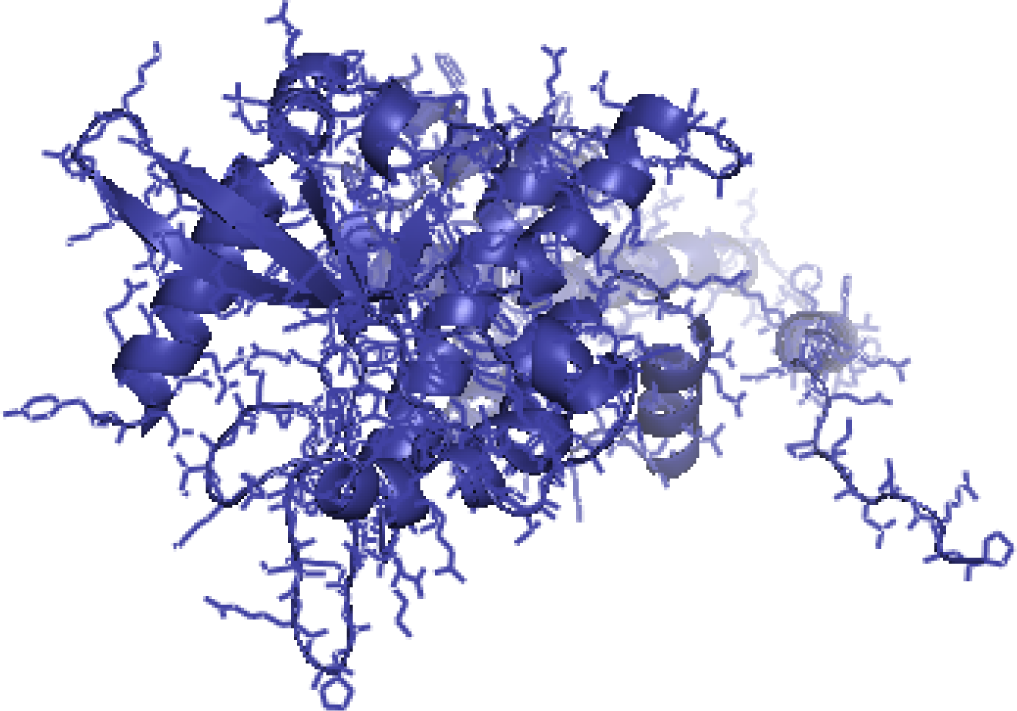
3D structure for STING for Duck.

Functional domains identified for the Domain in Apoptosis and Interferon response (DAPIN) is an 80-100 residue domain found at the N-terminus of various vertebrate-specific viral proteins involved in apoptosis, cancer, inflammation and immune response. The DAPIN domain can be found alone or in association with other domains such as CARD, LRR, SPRY, Caspase or zinc finger B-box. It has been proposed that the DAPIN domain could have adapter function, coupling apoptosis and immune disorders. The DAPIN domain has been shown to be a protein-protein interaction domain capable of binding to other DAPIN domains. Secondary structure predictions identified the DAPIN domain as predominantly α-helical, and it was suggested that it might belong to a superfamily of composite DEATH domains that includes a CARD, a DEATH domain (DD), and a DEATH effector domain (DED).

The DAPIN domain has also been called pyrin domain, pyrin N-terminal homology domain (PYD) or PAAD (after pyrin, AIM, ASC death-domain-like protein families). The NACHT domain is an evolutionarily conserved protein domain. This NTPase domain is found in apoptosis proteins as well as proteins involved in MHC transcription.[1] Its name reflects some of the proteins that contain it: NAIP (NLP family inhibitor of apoptosis protein), CIITA (i.e., C2TA or MHC class II transcriptional activator), HET-E (incompatibility locus protein from Podospora anserina), and TEP1 (i.e., TP1 or telomerase-associated protein).

The NACHT domain contains 300 to 400 amino acids. It is a predicted nucleoside triphosphatase (NTPase) domain found in animal, fungal and bacterial proteins. It is found in association with other domains such as a CARD domain (InterPro: IPR001315), a pyrin domain (InterPro: IPR004020), a HEAT repeat domain (InterPro: IPR004155), a WD40 repeat (InterPro: IPR001680), a leucine-rich repeat (LRR) or BIR repeats (InterPro: IPR001370).

The NACHT domain consists of seven distinct conserved motifs, including an ATP/GTPase-specific P-loop, an Mg 2+-binding site (Walker motifs A and B, respectively), and five more specific motifs. Unique features of the NACHT domain include the prevalence of “minor” residues (glycine, alanine, or serine) directly C-terminal to the Mg 2+-coordinating aspartate in the Walker B motif, instead of the second acidic residue prevalent in other NTPases. The second acidic residue is typically found in NACHT-containing proteins two positions downstream. In addition, distal motif VII contains a conserved pattern of polar, aromatic, and hydrophobic residues not seen in any other NTPase family.

### Differential mRNA expression profiling for Mitochondrial genes

Differential mRNA expression profiling for cytochrome B and cytochrome C have been depicted in Fig 6 and Fig 7 respectively. Gene expression of both Cytochrome B and cytochrome C was observed to be better in healthy ducks in comparison to infected ducks.

**Fig 6:**
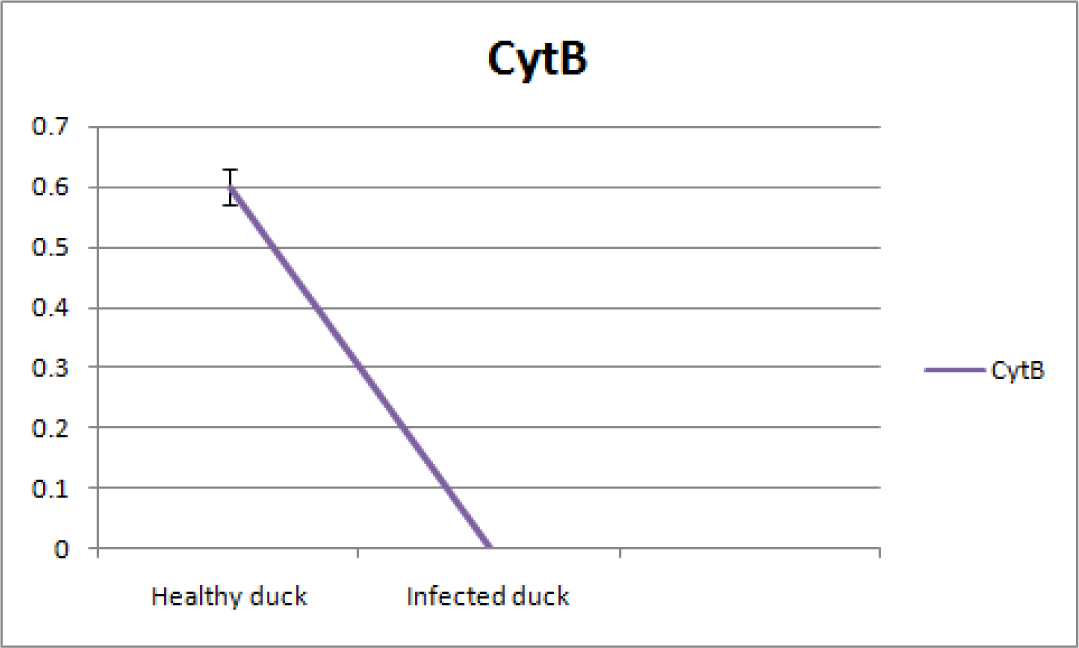
Differential mRNA expression profiling for Cytochrome B gene in Duck with respect to *Pasteurella multocida* infection

**Fig 7:**
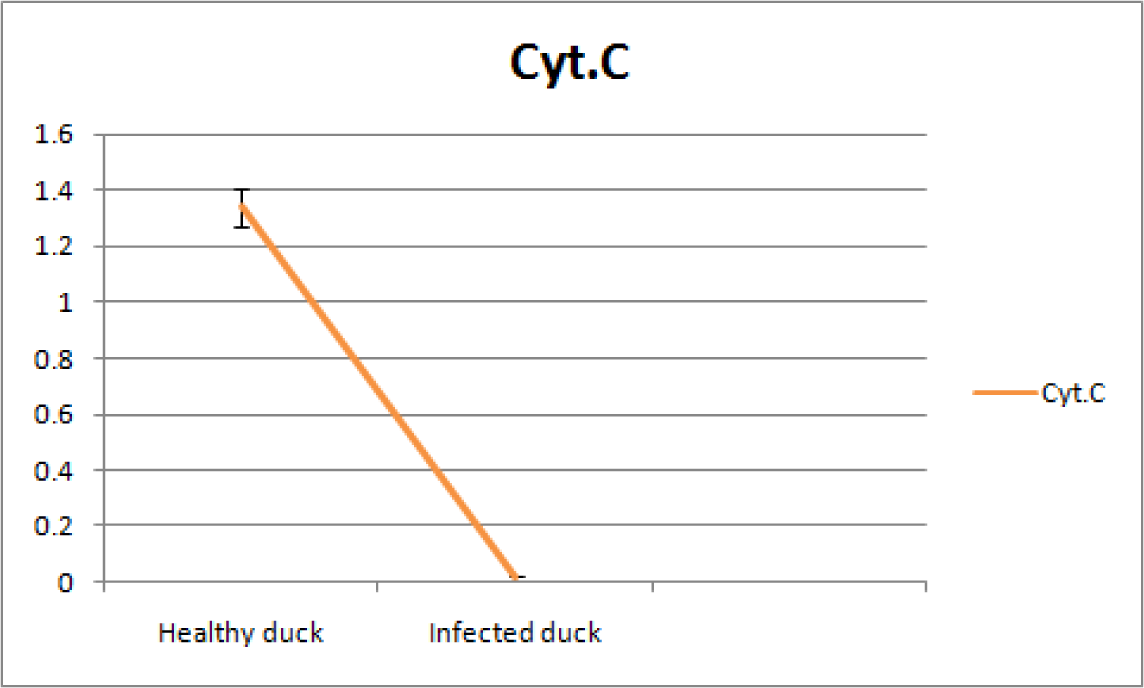
Differential mRNA expression profiling for Cytochrome C gene in Duck with respect to *Pasteurella multocida* infection

**Fig 8:**
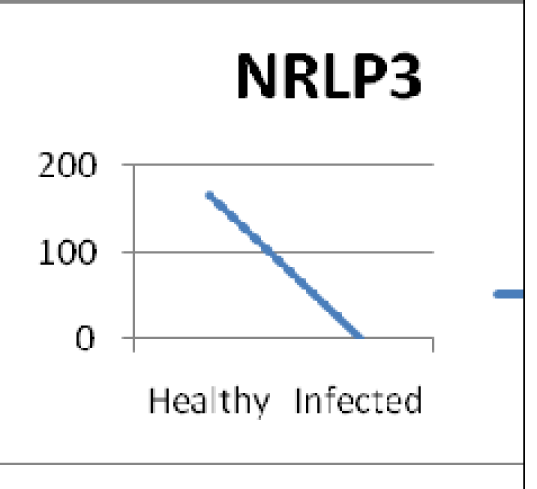
Differential mRNA expression profiling for NRLP3 gene in duck

**Fig 9:**
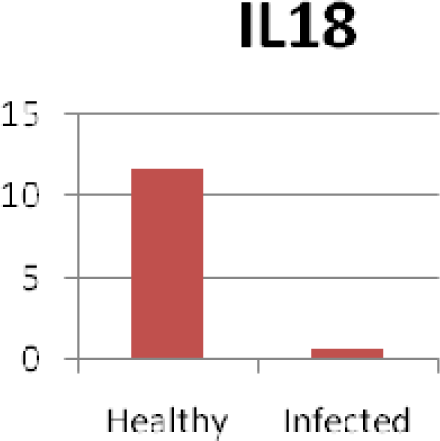
Differential mRNA expression profiling for IL18 gene in duck

### Differential mRNA expression profiling for nuclear(NRLP3, IL18 and STING) genes

Differential mRNA expression profiling for NRLP3, IL18 and STING gene in duck liver tissue reveal better expression of these genes in helathyducks in comparison to infected ducks.

### Molecular docking of IL18 with Pasteurella multocida

Molecular docking analysis was conducted for IL18 with certain surface protein of P*asteurella multocida* predicted to bind with membrane protein, lipoprotein, zinc transporter membrane, electron transporter RnfD, cell division protein FtsW.

Patchdock score was observed to be high (21096) for IL18 with Pasteurella multocida membrane protein. On the otherhand patchdock score was less for lipoprotein 14344. And zinc transporter membrane as 14598, lipopolysaccharide16374, <u>Pasteurella_multocida</u> <u>electron_transporter_RnfD</u> 19176.,cell division protein FtsW 16392.

Molecular docking for IL18 with the duck pasteurellamultocida membrane protein indicates a good score and is represented in Fig 11a. (IL18: Blue for IL18, red for membrane protein), whereas Fig 11b indicates the binding sites for IL18 with the *Pasteurella multocida* membrane protein.

**Fig 10:**
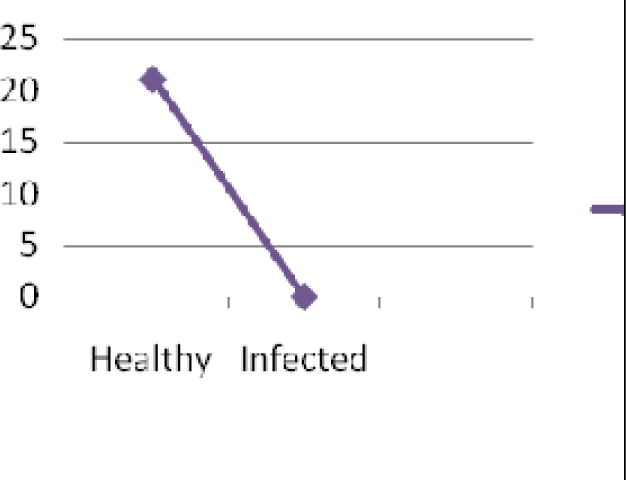
Differential mRNA exression profiling for Sting gene w.r.t Pasteurellosis in liver tissue.

**Fig 11A.**
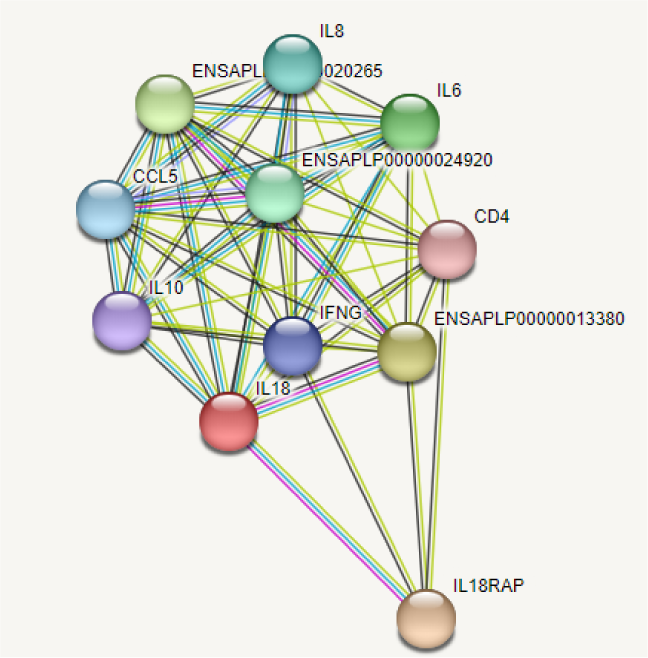
Molecular interaction analysis for IL18 with other genes.

**Fig 11B.**
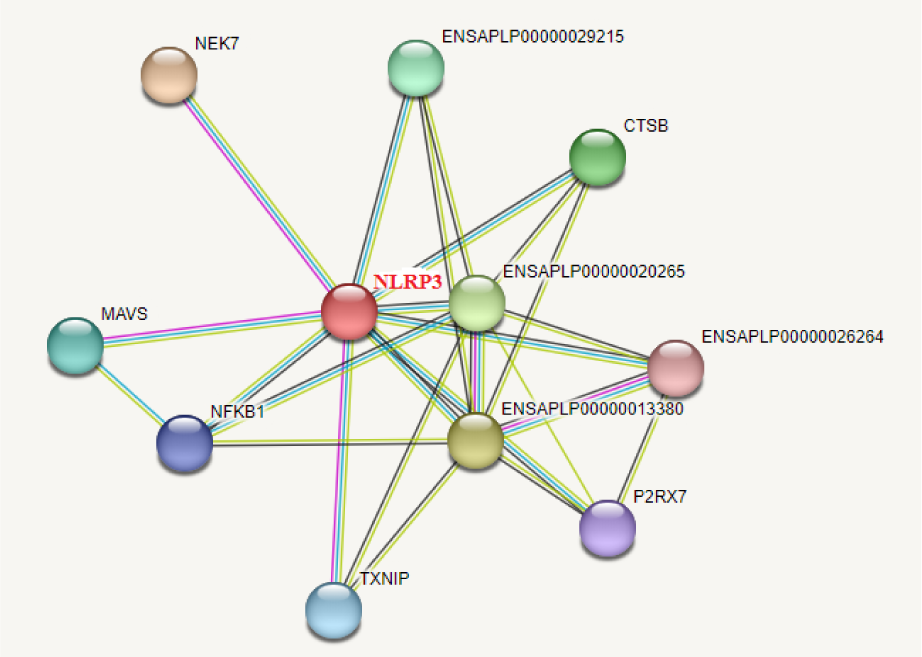
Molecular interaction analysis for NRLP3 for duck with other genes.

**Fig 11C.**
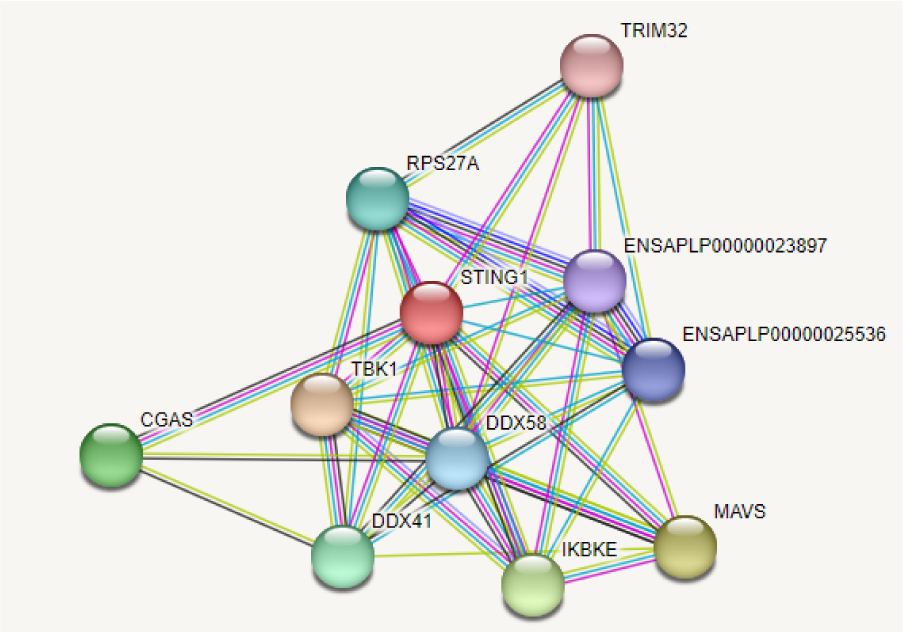
Molecular interaction analysis for STING for duck with other genes.

### Interaction String Analysis

In any biological process, the proteins act together and interact with each other. The molecular interaction of IL18, NRLP3and STING protein have beendepicted in Fig 11a, Fig 11b and Fig 11c.

### Kegg Analysis

The biological pathway analysis for IL10, NRLP3 genes are being depicted in Fig 12 A to D.

**Fig 12A:**
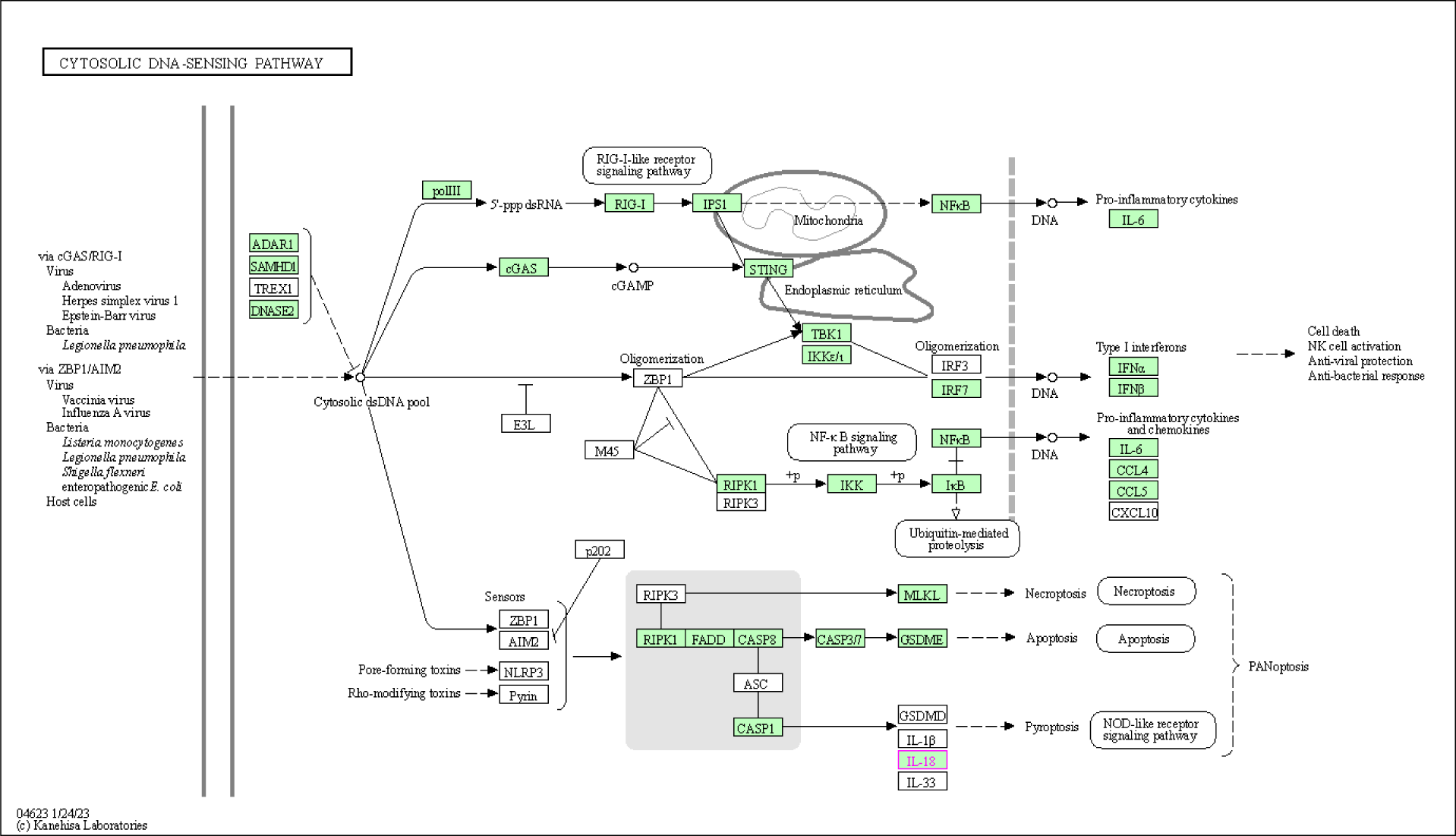
Molecular pathway analysis for IL18 gene in duck.

**Fig 12B:**
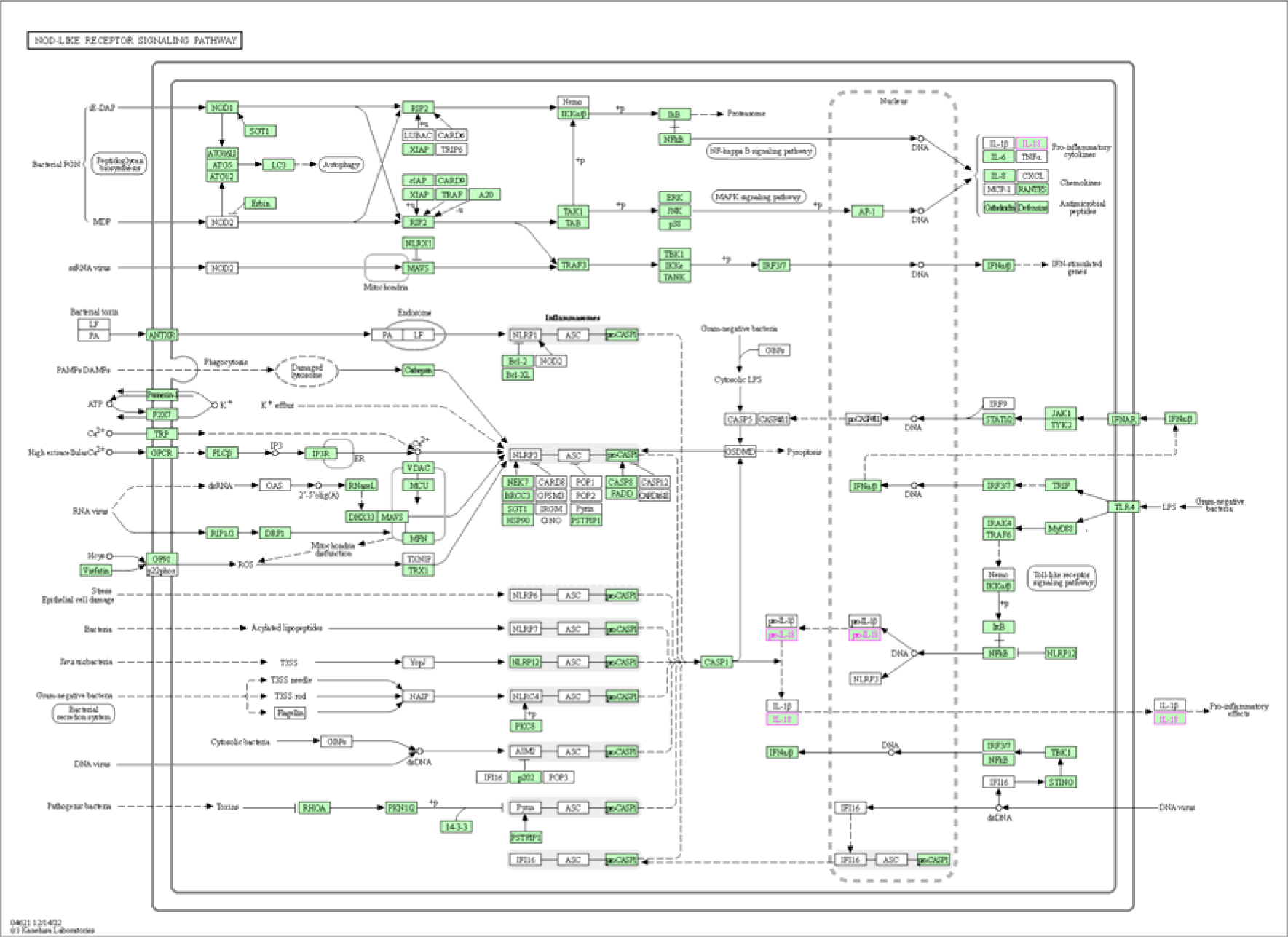
Molecular pathway analysis for IL18 and NRLP3 gene in duck.

**Fig 12C:**
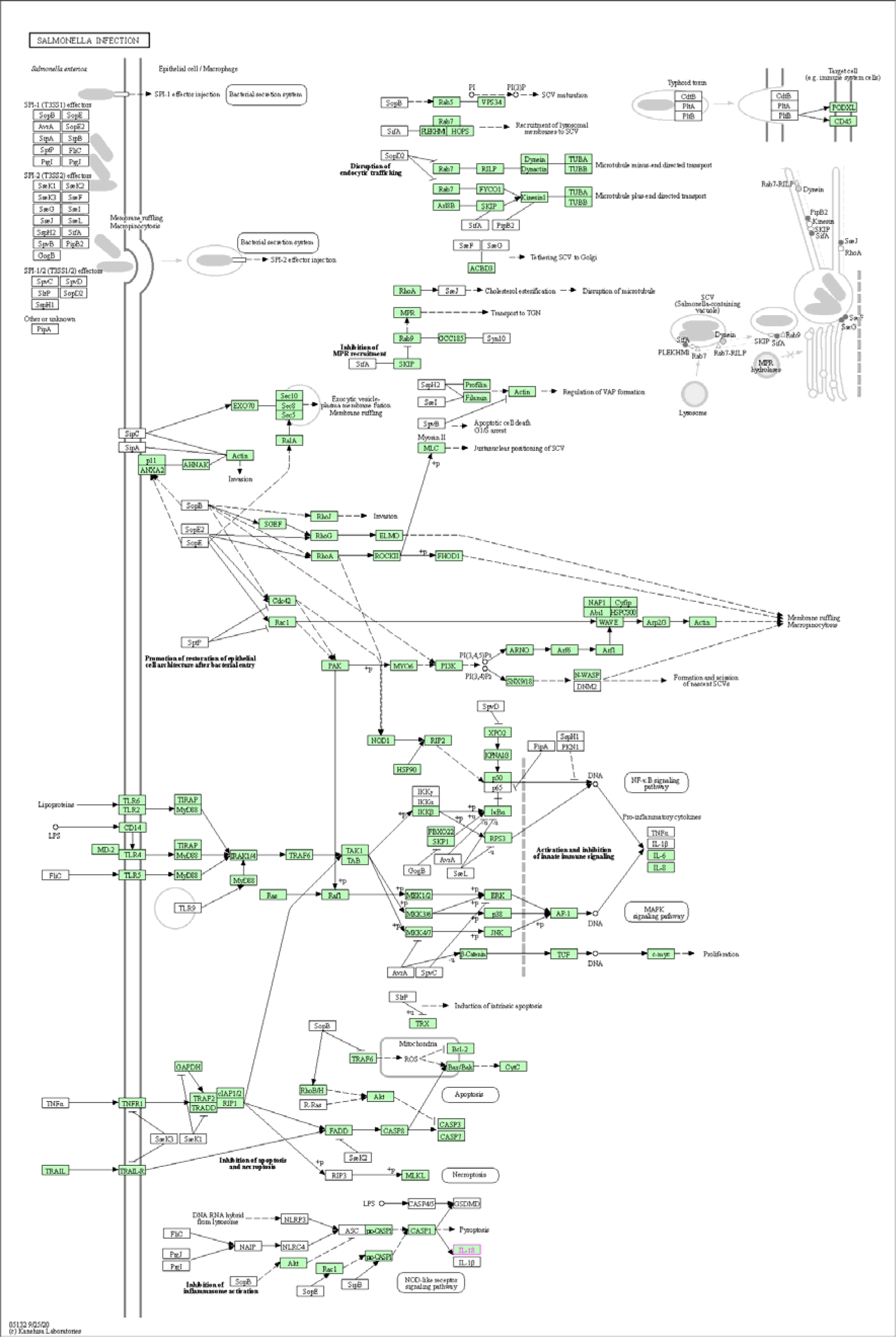
Molecular pathway analysis for IL18 gene in duck in response to bacterial infection.

**Fig 12D:**
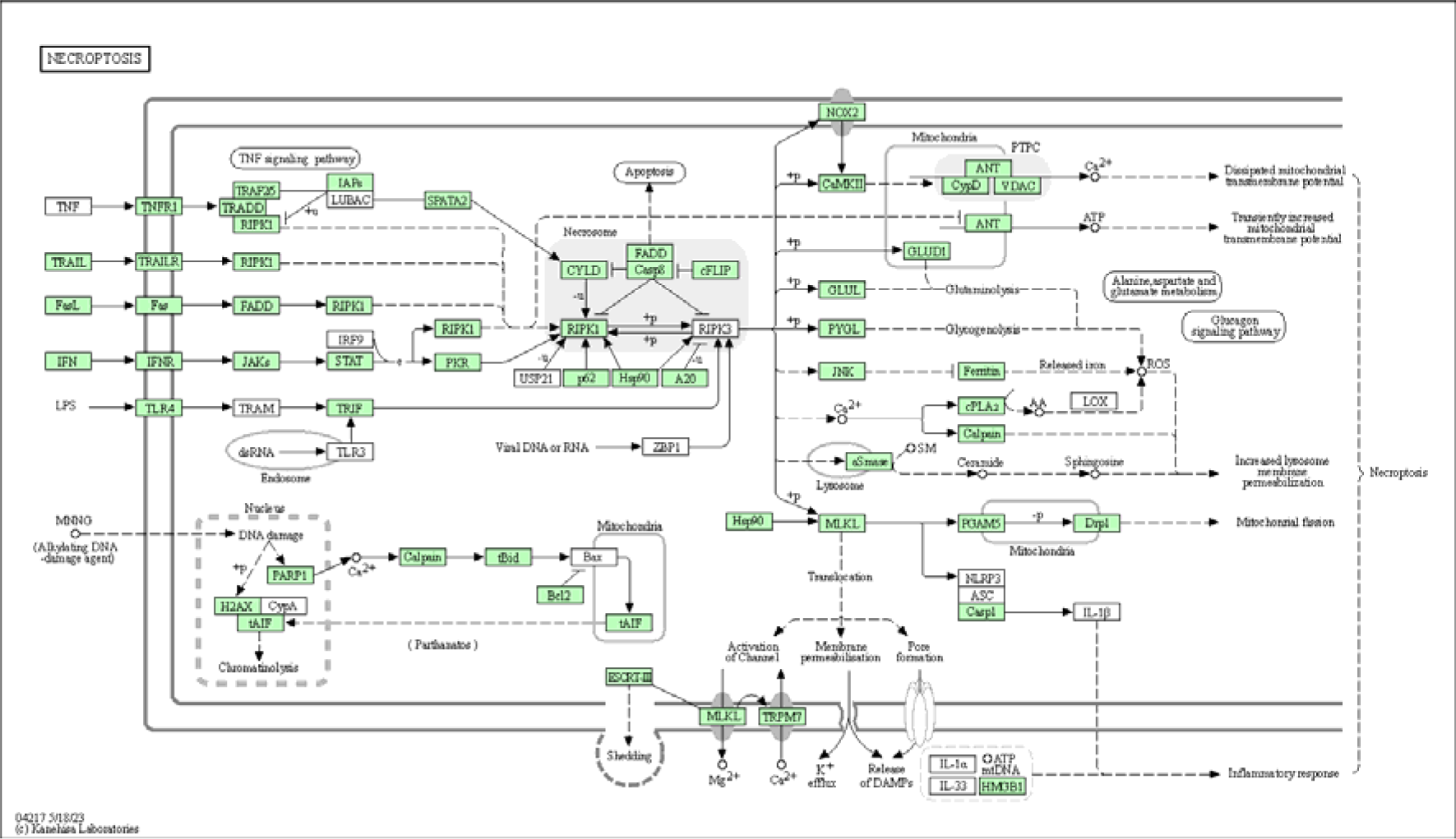
Molecular pathway analysis for NRLP3 gene and interaction of mitochondria DNA with nuclear genes in duck.

## Discussion

In this current research, we attempt to explore the molecular control of host resistance against duck Pasteurellosis in duck model. We attempt to identify the role of mitochodrial gene and its role against duck Pasteurellosis. We have also tried to explore the interaction of mitochondrial gene with nuclear gene against bacterial infection. This is a novel finding and first report of this kind.

We have characterized and identified the important domains for two polypepide coding mitochondrial genes, namely cytochrome B and Cytochrome C for indigenous Bengal Duck. We have also characterized immune response genes for Bengal Duck, namely IL18, NRLP3 and STING. In the next step, the experimental ducks were naturally infected with *Pasteurella multocida*. Some ducks were not affected, we regard them as *healthy group*, while some ducks were infected and died later on, and we regarded them as *infected group*. We have repeated the experiment *in ovo* with the isolated Pathogenic *Pasteurella spp.* We collected liver and kidney samples from both the groups and subjected to differential mRNA expression profiling. It is to be noted that we have isolated mRNA for mitochondrial genes without isolating the mitochondria. We speculate that the mRNA isolated were from cytoplasm, i.e. mRNA transcribed from released or escaped mitochondrial DNA from the mitochondria.

We observed better expression for cytochrome B and cytochrome C in the healthy ducks in comparison to that of infected ducks. We have also observed better expression of IL18, NRLP3 and STING genes in healthy ducks in comparison to infected group. Hence we observed positive correlation of mitochondrial gene expression along with the identified nuclear genes. KEGG analysis have revealed the role of mtDNA escape from the mitochondria, which triggers various pathways including Cytosolic DNA sensing pathway, NOD like signalling receptor pathway, pathway depicting howthebody’s immune response is triggered with Salmonella infection, necroptosis pahway. STRING analysis have revealed that how various immune response genes interact with each other and acts with CASCADE formation.

Earlier studies reported that mtDNA escapes during infection and triggers nuclear-derived immune response genes and then acts against and destroys the invading pathogen in a variety of ways^42^. Healthy ducks that survived exposure are expected to have a better mtDNA-mediated and triggered immune response. mtDNA released from mitochondria is able to activate cGAS-STING signaling. Researchers have examined mtDNA release in the context of cell death^43,44^, while West et al provided evidence that TFAM deficiency promotes mitochondrial stress and mispackaged mtDNA^45^, leading to their expulsion into the cytoplasm where they bind and activate cGAS and initiate the type I interferon response^46^.

Infection with the bacterial pathogen Mycobacterium tuberculosis triggers cGAS activation and subsequent IRF3^47–49^-dependent type I interferon response. This was thought to be due solely to the detection of mycobacterial DNA, but other studies have identified a role for mitochondrial stress and the subsequent release of mtDNA into the cytoplasm^50^. This observation is strain-dependent, but suggests a role for mitochondrial stress and dynamics in M. tuberculosis-induced mtDNA release. Previous work observed cytochrome C release from mitochondria in M. tuberculosis infected cells, suggesting that there may be a possible role for BAX/BAK51-dependent mitochondrial permeabilization in infection-associated mtDNA release^51^. Several researchers have depicted the role of mitochondrial DNA in cell death, apoptosis and their possible influence on immunity or disease resistance^52–55^. Mitochondrial permeability is dependant on nuclear factor NF-κβ-dependent anti-tumour activity under caspase deficiency^56^. The role of cytochrome C in maintaining the integrity of mitochondria and its association with disease has been reported by some workers^57–60^. The nuclear DNA is ready to be expelled as a NET through a highly regulated process involving chromatin decondensation and citrullination histones. Furthermore, permeabilization of the plasma membrane it is also regulated, which inevitably leads to cell death^61^. The complex cross talk between mitochondria and NRLP3^62, 63^ and NALP3^64^ has been studied. The complex interaction of mitochondrial permeability and influx of mitochondrial DNA have been reported^65–67^ with their possible role with disease incidences, like myocardial infarction^68^. Role of mitochondrial genes in controlling innate immunity against bacterial infection has been reported^69,70^.Hence it is evident that there exists a complex interaction between mitochondrial and nuclear gene and its association with diseases.

## Acknowledgement

The authors are thankful to Department of Biotechnology, Ministry of Science and Technology, Govt. of India (Grant number BT/PR24310/NER/95/649/2017) and Department of Science and Technology, Govt. of India (Grant no. EMR/2016/003554) for providing the financial support. The technical and financial support by Vice-Chancellor, West Bengal University of Animal and Fishery Sciences is duly acknowledged. Thanks to Director, AH & VS, Animal Resource Development Department, Govt. of West Bengal. Authors are thankful to Xcerlis Lab Ltd. for provision of NGS instrumentation facilities for conducting the metagenomics work.

